# Discovery of new deaminase functions by structure-based protein clustering

**DOI:** 10.1101/2023.05.21.541555

**Authors:** Jiaying Huang, Qiupeng Lin, Hongyuan Fei, Zixin He, Hu Xu, Yunjia Li, Kunli Qu, Peng Han, Qiang Gao, Boshu Li, Guanwen Liu, Lixiao Zhang, Jiacheng Hu, Rui Zhang, Erwei Zuo, Yonglun Luo, Yidong Ran, Jin-Long Qiu, Kevin Tianmeng Zhao, Caixia Gao

## Abstract

The elucidation of protein function and its exploitation in bioengineering have greatly contributed to the development of the life sciences. Existing protein mining efforts generally rely on amino acid sequences rather than protein structures due to technical difficulties in structural elucidation. We describe here for the use of AlphaFold2 to predict and subsequently cluster an entire protein family based on predicted structure similarities. We selected the deaminase family of proteins to analyze and through this approach identified many previously unknown deaminase properties. We applied these new deaminases to the development of new cytosine base editors with distinct features. Although we found many new double-stranded DNA deaminases from the DddA-like protein clade, we were surprised to find that most of the proteins in this family were not actually double-stranded DNA cytidine deaminases. From this protein clade, we engineered the smallest single-strand specific cytidine deaminase, which facilitates the first efficient cytosine base editor to be packaged into a single AAV vector. Importantly, we also profiled a deaminase from this clade that edits robustly in soybean plants, which previously suffered from poor editing by cytosine base editors. These newly discovered deaminases based on AI-assisted structural predictions greatly expand the utility of base editors for therapeutic and agricultural applications.

## Introduction

The discovery and engineering of new proteins has greatly transformed the life sciences. Traditional enzyme mining based solely on sequence information has been effective at classifying and predicting protein functions and evolutionary trajectory^1, 2^. However, one-dimensional (1D) information, whether in the form of core amino acids, specific motifs, overall amino acid sequence identity, or Hidden Markov Models (HMM), cannot completely illuminate the functional characteristics of proteins.

In contrast, since protein function is ultimately determined by three dimensional (3D) protein folds, understanding protein structures would provide reliable and rational insights into protein function during the process of protein mining and clustering classifications^3, 4^. Although the number of publicly reported protein structures is increasing, it is miniscule compared to the number of new proteins discovered based on amino acid sequences^5, 6^. Recently, many artificial intelligence (AI) methods have been developed that use 1D amino acid sequences to accurately predict high resolution 3D protein structures^7–9^. These protein structure prediction methods should thus enable large-scale mining and classifications of proteins with specific functions.

Deaminase-like proteins catalyze the deamination of nucleotides and bases in nucleic acids. They play important roles in defense, mutation and nucleic acid metabolism and other biological processes^10–13^ and have been recently exploited for use in programmable DNA and RNA base editors^14–16^, a class of precise genome editing technologies. Members of this family act as nucleotide deaminases and nucleic acid deaminases, including adenosine, cytidine, cytosine and guanine deaminases, and have the ability to act on single-stranded DNA (ssDNA)^17^, double-stranded DNA (dsDNA)^10^, double-stranded RNA (dsRNA)^18^, transfer RNA (tRNA)^19^, free nucleosides^12^, and other deaminated nucleotide derivatives^20^. The sporadic distribution of deaminases and their rapid evolution due to positive selection often confounds the relationships between the various protein families in phylogenetic analyses based on sequence^20, 21^. Here, we performed new protein clustering classifications on the greater deaminase family of proteins based on AlphaFold2-predicted 3D structures.

To better differentiate and discover deaminases with diverse functions, we employed AlphaFold2 to first predict deaminase structures and subsequently performed structural comparisons to generate a new taxonomic tree of deaminase proteins that better reflect the different types of cytidine deaminases. Using AlphaFold2-predicted structures, we were able to classify proteins into different clades more efficiently than using 1D amino acid sequences.

Cytosine base editors (CBEs) use cytidine deaminases to catalyze C-to-U base conversions, resulting in permanent C • G-to-T • A base edits in DNA^14, 15, 22, 23^. Base editors have great potential in therapeutic genome editing, fundamental life sciences research, and for breeding new elite traits into plants^24–26^. Previous DNA base editors exploited the use of two types of cytidine deaminases acting on either ssDNA or dsDNA^10, 14^. To date, only a few ssDNA-targeting APOBEC/AID-like deaminases and one dsDNA-targeting deaminase (DddA) have been used to generate CBEs^10, 14, 15, 27–30^. These deaminases remain limited to sequence context restrictions, low on-off target editing ratios and large protein sizes, which makes their delivery by adeno-associated virus (AAV) viral vectors difficult^31^. For unknown reasons, some species like soybean plants, a staple agricultural crop grown all over the world, have suffered from poor cytosine base editing since the technology was first introduced in 2016^32^. Thus, robust and more efficient CBEs are still needed to further expand their utility. By generating new protein classifications based on their predicted structures, we have developed a suite of new ssDNA and dsDNA deaminases used for precision genome editing. We highlight that enzyme mining based on structures predicted by AlphaFold2 is a simple, flexible, and high-throughput method to classify and engineer proteins with unknown functions.

## Results

### Clustering and discovery of new cytidine deaminases via protein structures

We hypothesized that the comparison and clustering of known or predicted protein structures, given that the 3D structure of a protein ultimately determines its function, could be an effective method for classifying deaminases into functional clades. Thus, we employed a combination of AI-assisted protein structure prediction, structural alignments, and clustering to generate new protein classification relationships among deaminases (Figure 1A). We selected 238 protein sequences annotated as having a deaminase domain from the InterPro database and 4 distant outgroup candidate protein sequences from the JAB-domain family (Figure S1A). Specifically, we randomly selected 15 candidates of at least 100 amino acids in length from each of the 16 deaminase families and used AlphaFold2 to predict their protein structures. We conducted multiple structural alignments (MSTA) of all candidates using normalized TM-scores as a guide^33^. Based on the MSTA results, we generated candidate similarity matrices reflecting the overall structural correlation between the proteins. We then organized these similarity matrices into a structural dendrogram using the average-linkage clustering algorithm (Figure 1B). The dendrogram clustered the 238 proteins into 20 unique structural clades and the deaminases within each clade had distinct conserved protein structural domains (Figure 1C and 1D).

**Figure 1.**
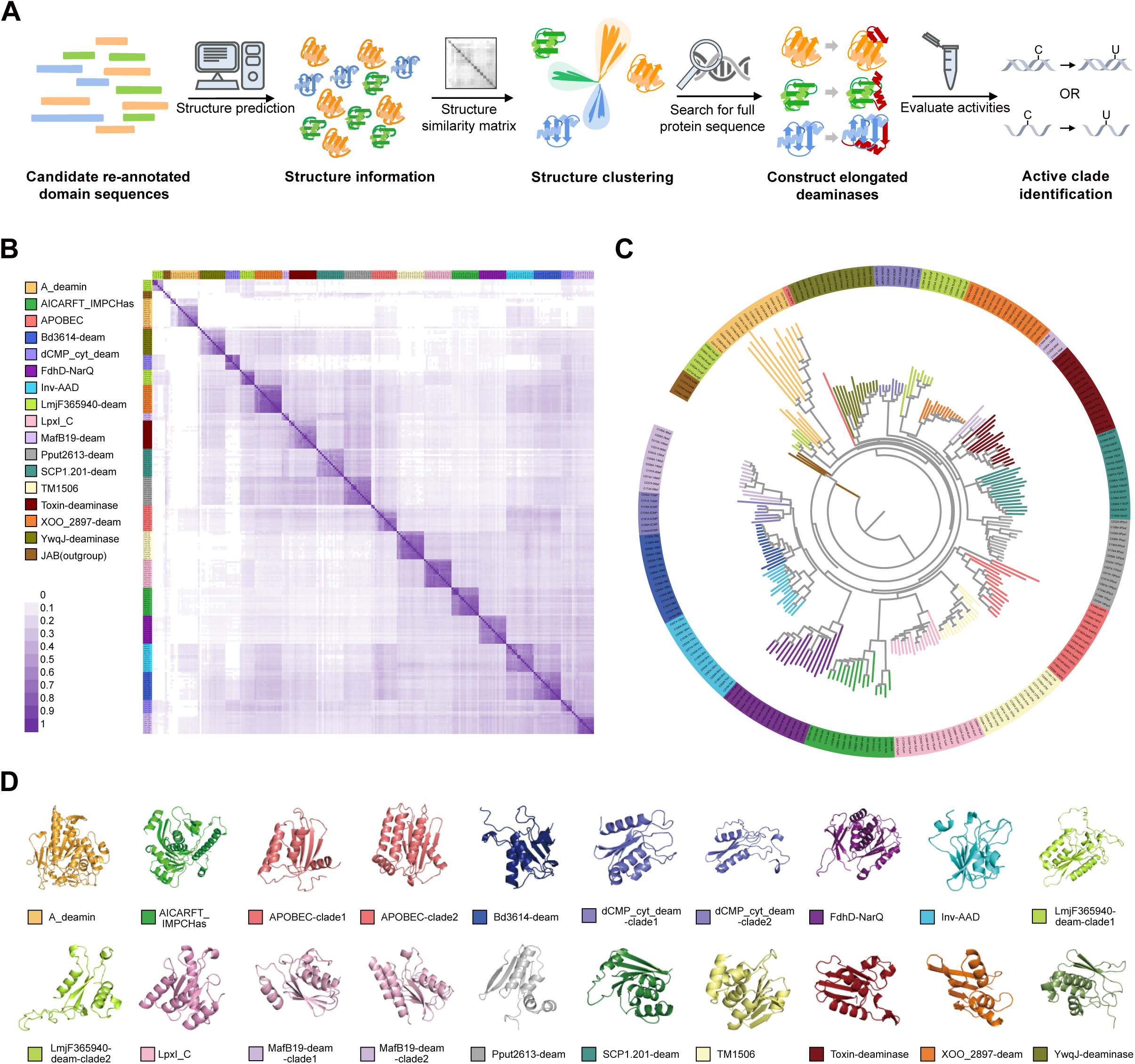
Protein clustering of deaminases based on structures predicted by AlphaFold2. (A) Workflow of protein clustering based on AlphaFold2-predicted structures. The structures of candidate re-annotated domain sequences were predicted by AlphaFold2 and subsequently clustered based on structural similarities. Then, ssDNA and dsDNA cytidine deamination activities were experimentally tested in plant and human cells. (B) Structural similarity matrix to reflect similarities between 242 predicted protein (238 cytidine deaminases and 4 JAB) structures across 16 deaminase families and one outgroup. Different family proteins are distinguished by different colors; heat map color shades indicate the degree of similarity. (C) The classification of proteins into different deaminase families based on protein structure and labeled with different color modes. (D) Representative predicted structures for each of 16 deaminase clades.

We found that accurate protein clustering classifications could be generated base on protein structural alignments, even without the use of contextual information such as conserved gene neighborhoods and domain architectures. When using structure-based hierarchical clustering, different clades reflected unique structures, implying distinct catalytic functions and properties (Figure 1D). Interestingly, we also found that this structure-based clustering method was much more effective at sorting for functional similarities than traditional 1D amino acid sequence-based clustering approaches. For example, adenosine deaminases (A_deamin, PF02137 in InterPro database), enzymes involved in purine metabolism, were split into different clades when using amino acid sequence-based clustering methods but were all grouped together into a single A_deamin-clade using our structure-based clustering approach (Figure 1B, 1C and S1B). Additionally, four deaminase families (dCMP, MafB19, LmjF365940 and APOBEC as annotated by InterPro) were each divided into two separate clades when using structure-based clustering (Figure 1C and 1D). Comparison of protein structures showed that the two clades for each of these four deaminase families had quite different structures, contrary to what their InterPro naming and sequenced-based classification might suggest (Figure 1D and S1C). In summary, AI-assisted 3D protein structures provide reliable clustering results and only require an amino acid sequence from the user, making it a convenient and effective strategy for generating protein relationships.

### Evaluating diverse deaminase clades by fluorescence imaging

CRISPR-based CBEs are precise genome editing technologies capable of generating C•G-to-T•A substitutions in the genome of living cells. Because single-strand DNA specific cytidine deaminases are an essential component of CBEs, we sought to explore the deamination activity of each structure-based classified deaminase clade in the context of DNA base editing. We evaluated a total of 190 deaminase domains by selecting at least five proteins from each clade. Importantly, because the core deaminase domain used for clustering may not show editing activity, we extended each deaminase sequence to include additional secondary structures from each corresponding gene around the deaminase domain (Figure S1A). For each of 190 newly annotated deaminases, we generated plant CBEs by fusing each candidate domain-related sequence to the N-terminus of a Cas9 nickase (nCas9, D10A) followed by an uracil-DNA glycosylase inhibitor (UGI)^14, 34^. We developed four BFP-to-GFP reporter systems to reflect T<u>C</u>, C<u>C</u>, G<u>C</u>, and A<u>C</u> 5’-base deamination preferences (Figure S2A). Each CBE was co-transformed with all four BFP-to-GFP reporter plasmids into rice protoplasts and analyzed by fluorescent microscopy after three days^34^. We found that deaminases belonging to the SCP1.201 (PF14428), XOO_2897 (PF14440), MafB19 (PF14437), Toxin-deaminase (PF14424), and TM1506 (PF08973) clades possessed ssDNA cytidine deamination activity. Interestingly, we noticed that some deaminase candidates displayed different sequence preferences compared to the APOBEC/AID-like deaminases as evaluated using the fluorescence reporter system. Therefore, we demonstrated that the use of 3D structures for protein classification enabled the discovery of new functional deaminase clusters for use in base editors, offering new opportunities for developing enhanced and bespoke precise base editing tools.

### Validation of the diverse functions of SCP1.201 deaminases

While evaluating deaminases from each clade, we were surprised to find that some deaminases annotated from the SCP1.201 clade were capable of deaminating single-stranded DNA substrates. These deaminases were previously named <u>D</u>ouble-stranded <u>D</u>NA <u>d</u>eaminase toxin <u>A</u>-like (DddA-like) deaminases in the InterPro database (PF14428). The DddA deaminase was recently developed into a CRISPR-free double-stranded DNA cytosine base editor (DdCBE) capable of deaminating cytosine bases on double-stranded DNA^10^. Because of DddA, all proteins in the SCP1.201 clade were also annotated as double-stranded DNA deaminases. To re-analyze this SCP1.201 clade, we selected all 489 SCP1.201 deaminases from the InterPro database. We also included seven additional proteins that were 35% to 50% identical by Basic Local Alignment Search Tool (BLAST) with DddA but were characterized separately in InterPro. After identity and coverage filtering, we performed a new AI-assisted protein structure-based classification of 332 SCP1.201 deaminases. Structure clustering showed that the SCP1.201 deaminases clustered into different clades with unique core structural motifs (Figure 2A-2E).

**Figure 2.**
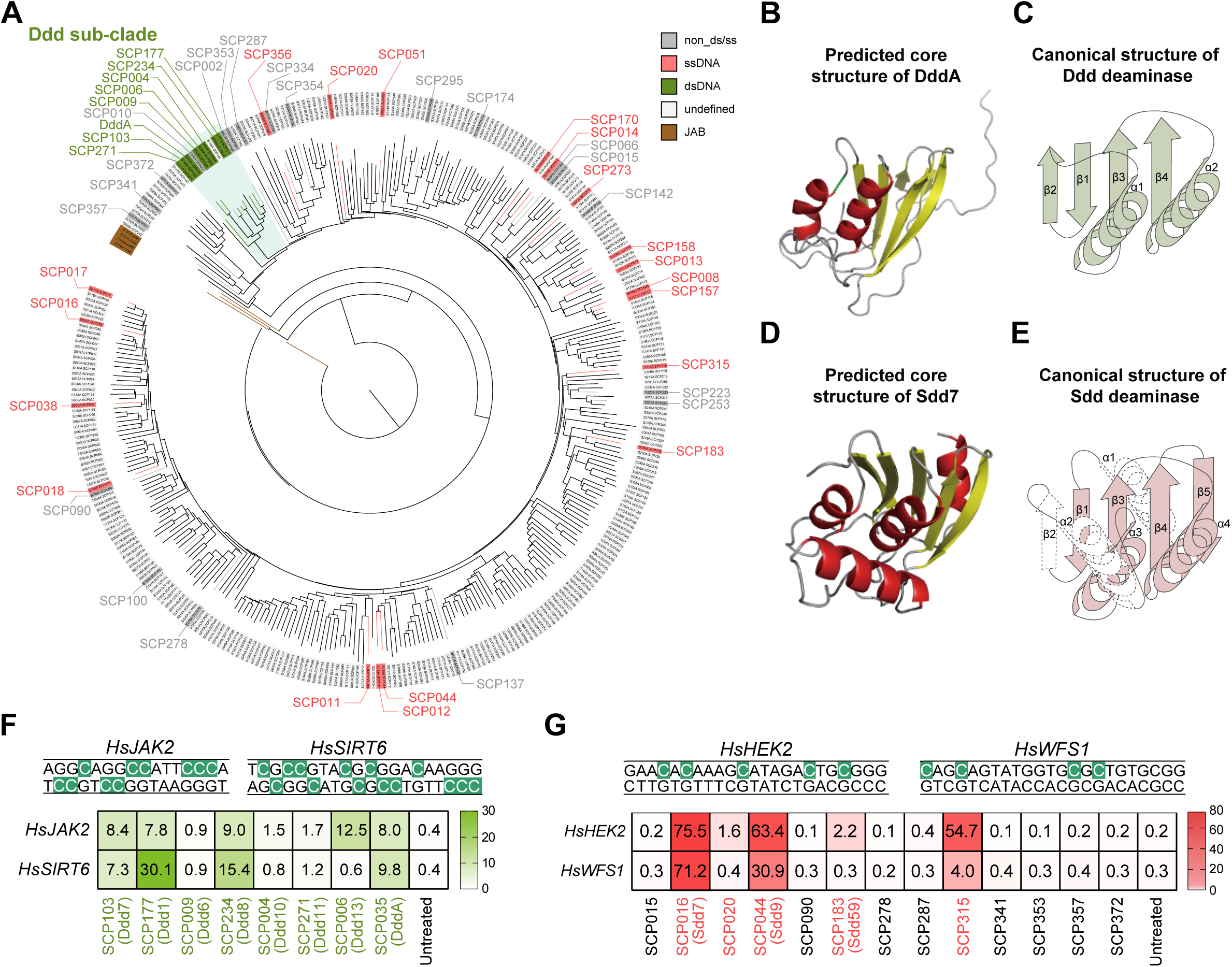
The clustering and characteristics of SCP1.201 deaminases. (A) Classification of SCP1.201 deaminases based on protein structure. The JAB families are colored brown and regarded as an outgroup, and the tested deaminases are shown in red (single-strand editing), green (double-strand editing) or dark grey (no editing). Undefined deaminases in light grey await further functional analysis. (B) Predicted core structure of DddA by AlphaFold2. (C) Characteristics of the canonical structure of Ddd protein. (D) Predicted core structure of Sdd7 by AlphaFold2. (E) Characteristics of the canonical structure of Sdd protein. (F) Experimental evaluation of dsDNA deamination activity of Ddds at two endogenous sites in HEK293T cells. The edited bases used for calculating editing are highlighted in green. (G) Experimental evaluation of ssDNA deamination activity of Sdds at two endogenous sites in HEK293T cells. The edited bases used for calculating editing are highlighted in green. Data in (F) and (G) are representative of three independent biological replicates (*n* = 3).

We found that DddA and ten other proteins clustered into one subclade of SCP1.201. Upon analyzing the 3D predicted structures of all 11 proteins within this subclade, we found that they shared a similar core structure to DddA. Given their structural similarities to DddA, we hypothesized that the other proteins in this subclade can also perform double-strand DNA cytidine deamination. To evaluate dsDNA deamination, we generated DdCBEs comprised of each deaminase alone or split in half at a residue similar to the site where DddA was split by protein structure alignment and joined together using a dual TALE system^10^ (Figure S2B). We evaluated 10 proteins from this Ddd subclade in HEK293T cells at the *JAK2* and *SIRT6* sites and observed that 8 proteins could perform dsDNA base editing (Figure 2A and 2F). We hereafter named these deaminases as <u>d</u>ouble-strand <u>D</u>NA <u>d</u>eaminases (Ddd) and assigned them to this newly identified Ddd sub-clade.

To evaluate other SCP1.201 candidate proteins, we selected 24 proteins at random and subjected these to our CBE fluorescent reporter system. We found that 22 showed detectable fluorescence and selected 13 to evaluate endogenous base editing in the context of CBE in mammalian cells (Figure 2). Although these were previously characterized as DddA-like, many showed cytosine base editing activity on ssDNA (Figure 2A, 2G) but not dsDNA (Figure S2C). Therefore, we hereafter named these ssDNA-targeting protein domains from the SCP1.201 clade as single-stranded DNA deaminases (Sdd). We were surprised to find that a majority of protein members from the SCP1.201 clade were found to be Sdd proteins since these were all previously annotated as DddA-like. We also observed that these Sdd proteins shared a similar protein structure as Sdd7, one of the highest editing ssDNA CBEs, which is distinct from the Ddd proteins (Figure 2D and 2E). Thus, the annotated DddA-like deaminases in the InterPro database (PF14428) should be further subdivided and re-annotated accordingly.

In comparison, we also performed a clustering of the proteins from the SCP1.201 clade based on 1D amino acid sequences and found that some outgroup members were dispersed throughout the tree, even though we chose four more closely related families as outgroups (Figure S2D and S2E). These results highlight the usefulness and importance of using protein structure-based classifications for comparing and evaluating protein functional relationships.

### New Ddd proteins have distinct editing preferences to DddA

Due to the strict 5’-TC sequence motif preference of DddA, the use of DddA-based dsDNA base editors is limited predominantly to TC targets^10^. Although the recently evolved DddA11 displayed a broadened ability to deaminate and edit cytosine bases with a 5’-HC (H = A, C or T) motif, the editing efficiency for AC, CC, and GC targets still need to be improved^35^. We evaluated the newly discovered Ddd proteins to determine if they could expand the utility and targeting scope of DdCBEs. 13 deaminases belonging to the Ddd sub-clade were cloned into DdCBEs and evaluated for dsDNA base editing at the endogenous *JAK2* and *SIRT6* sites in HEK293T cells (Figure 2F, S3A, S3B). Interestingly, we found that Ddd1, Ddd7, Ddd8, and Ddd9 had comparable or higher editing efficiencies to DddA (Figure 3A, S3A and S3B). Importantly, we identified that Ddd1 and Ddd9 had a much higher editing activity compared to DddA at 5’-GC motifs (Figure 3A, S3A and S3B). Strikingly, at the C10 (5’-GC) residue in *JAK2* and the C11 (5’-GC) residue in *SIRT6*, we found that while DddA resulted in 21.1% and 0.6% editing, Ddd9 was capable of editing 65.7% and 45.7%, respectively (Figure 3A).

**Figure 3.**
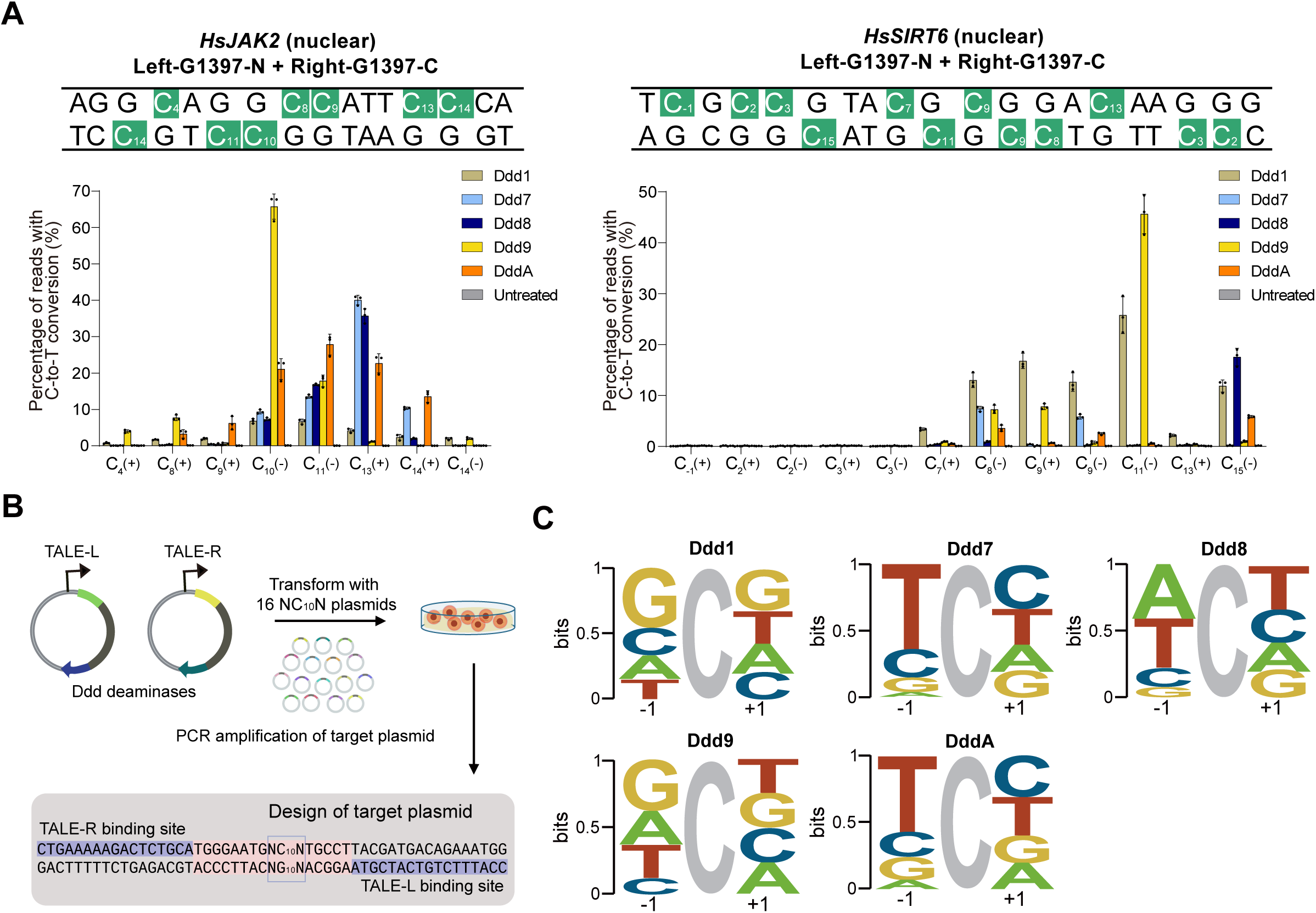
Evaluating newly discovered Ddd protein properties for use as base editors. (A) Editing efficiencies and editing windows of Ddd1, Ddd7, Ddd8, Ddd9 and DddA SCP1.201 dsDNA deaminases at two genomic target sites in HEK293T cells. (B) Plasmid library assay to profile context preferences of each Ddd protein in mammalian cells. Candidate proteins target and edit the “NC10N” motif. (C) Sequence motif logos summarizing the context preferences of Ddd1, Ddd7, Ddd8, Ddd9, and DddA as determined by the plasmid library assay. For all plots, dots represent individual biological replicates, bars represent mean values, and error bars represent the s.d. of three independent biological replicates (*n* = 3).

Because certain Ddd proteins seemed to exhibit distinct editing patterns compared to DddA, we sought to evaluate any sequence motif preference for these Ddd proteins. We first constructed 16 plasmids^35^ encoding the *JAK2* target sequence and modified positions 9-11 from G<u>C</u>C to N<u>C</u>N (N = A, T, C and G), yielding 16 different plasmids, and independently co-transfected each plasmid along with a DdCBE variant (Figure 3B). Following comparative analyses of C“ G-to-T ” A base conversion frequencies for each N<u>C</u>N, we generated corresponding sequence motif logos to reflect sequence context preferences of each dsDNA deaminase (Figure 3B). We found that as previously discussed, DddA and its structural homolog, Ddd7, strongly preferred a 5’-T<u>C</u> sequence motif (Figure 3C and S3C). In contrast, we found that Ddd1 and Ddd9 showed preferences towards editing 5’-G<u>C</u> substrates, while Ddd8 showed preferences towards editing 5’-W<u>C</u> (W=A or T) substrates. Therefore, the newly discovered dsDNA-targeting deaminases can edit cytosine bases at motifs previous inaccessible to DddA, which is also essential for future engineering efforts.

### Sdd deaminases enable base editing in human cells and plants

We next wondered whether the newly characterized Sdd proteins could be used for more precise or efficient base editing. We chose to evaluate the six most active Sdds as well as four weaker Sdds and compared their activities using a fluorescent reporter system. We generated plant CBEs for each of the ten Sdds and evaluated their endogenous base editing across six sites in rice protoplasts (Figure 4A and S4A). We found that seven of the deaminases (Sdd7, Sdd9, Sdd5, Sdd6, Sdd4, Sdd76 and Sdd10) had higher activity compared to the rat APOBEC1 (rAPOBEC1)-based CBE. The most active Sdd7 base editor reached as high as 55.6% cytosine base editing, which was more than 3.5-fold that of rAPOBEC1. To examine the versatility of these deaminases, we also constructed the corresponding human-cell targeting BE4max vectors^36^ and evaluated their editing efficiencies across three endogenous target sites in HEK293T cells. In agreement with the results in rice, we found that Sdd7 had the highest editing activity (Figure S4B).

**Figure 4.**
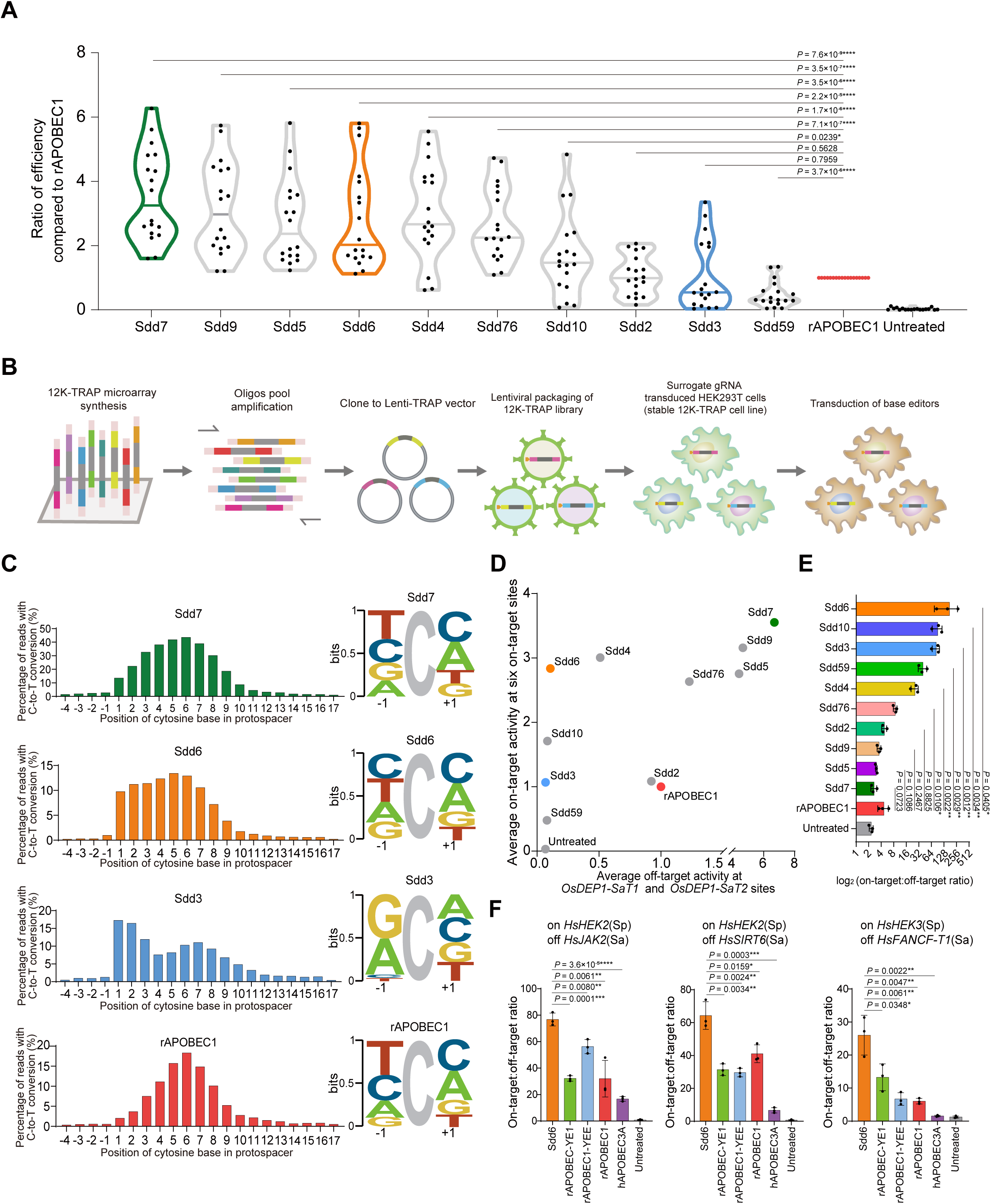
Evaluating newly discovered Sdd proteins for use as base editors in plant and human cells. (A) Overall editing efficiencies of the Sdds and rAPOBEC1 across six endogenous target sites in rice protoplasts. The average editing frequencies using rAPOBEC1 at each target were set to 1 and frequencies observed with Sdds were normalized accordingly. Dots represent each of three individual biological replicates across six endogenous genomic sites. (B) Overview of using 12K-TRAPseq to perform high throughput quantification of the activities and properties of the Sdds and rAPOBEC1 in HEK293T cells. (C) Overview of the editing properties and patterns of the Sdds and rAPOBEC1 as evaluated by the 12K-TRAP library. Left panels, the editing efficiencies and editing windows of the deaminases. Right panels, a sequence motif logo reflecting the context preferences of the deaminases. (D) Evaluation of off-target effects using an orthogonal R-loop assay in rice protoplasts. Dots represent average on-target C-to-T conversion frequencies of three independent biological replicates across six on-target sites in rice in (A) versus average sgRNA-independent off-target C-to-T conversion frequencies across two ssDNA regions (*OsDEP1-SaT1* and *OsDEP1-SaT2*) for each base editor. (E) On-target:off-target editing ratios for each base editor calculated from (D). (F) On-target:off-target editing ratios of Sdd6, rAPOBEC1-YE1, rAPOBEC1-YEE, rAPOBEC1, and hA3A tested across two on-target and three off-target sites in HEK293T cells. For (E) and (F), Dots represent individual biological replicates, bars represent mean values, and error bars represent the s.d. of three independent biological replicates (*n* = 3). Data are presented as mean values ± s.d. *P* values were obtained using two-sided Mann-Whitney tests. \**P* < 0.05, \*\**P* < 0.01, \*\*\**P* < 0.001, \*\*\*\**P* < 0.0001.

We previously showed that human APOBEC3A (hA3A) performed robust base editing with a large editing window in plants^37, 38^. We therefore compared the editing activities of hA3A and Sdd7 in human cells (Figure S4B) and plants (Figure S4C). Interestingly, Sdd7 had comparable editing activities as hA3A across all three target sites in HEK293T cells (Figure S4B) and five endogenous sites in rice protoplasts (Figure S4C). Because editing efficiency is of primary significance for genome editing in plant breeding, these results confirmed that Sdd7 is a robust cytosine base editor for use in both plants and human cells.

### Sdd proteins have unique base editing characteristics

When evaluating endogenous base editing, we observed different editing patterns by the different Sdd-CBEs across all tested genomic target sites in both human and rice cells. For instance, while Sdd7, Sdd9, and Sdd6 showed no particular motif editing preference, Sdd3 seemed to prefer editing 5’-G<u>C</u> and 5’-A<u>C</u> motifs and strongly disfavor editing 5’-T<u>C</u> and 5’-C<u>C</u> motifs (Figure S4D). To better profile the editing patterns of each deaminase, we used <u>T</u>argeted <u>R</u>eporter <u>A</u>nchored <u>P</u>ositional <u>Seq</u>uencing (TRAP-seq), a high-throughput approach for parallel quantification of base editing outcomes^39^. A 12K TRAP-seq library comprised of 12,000 TRAP constructs, each containing a unique gRNA expression cassette and the corresponding surrogate target site, was stably integrated into HEK293T cells by lentiviral transduction. Following cell culture and antibody selection, base editors were stably transfected into this 12K-TRAP cell line followed by ten days of blasticidin selection (Figure 4B). On the eleventh day post transfection, we extracted the genomic DNA and performed deep amplicon sequencing to evaluate the editing products of each deaminase (Figure 4B). We found that while Sdd7 and Sdd6 showed no strong sequence context preference, rAPOBEC1 had a strong preference for 5’-T<u>C</u> and 5’-C<u>C</u> bases while disfavoring 5’-G<u>C</u> and 5’-A<u>C</u> bases (Figure 4C). On the contrary, Sdd3 showed an entirely complementary pattern preferring to edit 5’-G<u>C</u> and 5’-A<u>C</u> bases while showing nearly no activity towards 5’-T<u>C</u> and 5’-C<u>C</u> bases (Figure 4C). Interestingly, we found that Sdd6 and Sdd3 had different editing windows and preferred to edit positions +1 to +3 in the protospacer as compared to rAPOBEC1 and Sdd7 (Figure 4C). In conclusion, the newly identified Sdd base editors show unique base editing properties such as increased editing efficiencies, disparate deamination motif preferences, and altered editing windows from conventional cytosine base editors.

It was previously described that CBEs could cause genome-wide Cas9-independent off-target editing outcomes, which raises concerns about the safety of these precise genome editing technologies for clinical applications^40, 41^. It is thought that these off-target mutations may be a result of overexpression of the cytidine deaminase. We wondered whether the newly-discovered Sdd proteins could offer a more favorable balance between off-target and on-target editing. We therefore evaluated the Cas9-independent off-target effects of the ten Sdds using an established orthogonal R-loop assay in rice protoplasts^42^. We found that six (Sdd2, Sdd3, Sdd4, Sdd6, Sdd10, and Sdd59) of the ten deaminases had lower off-target activities than rAPOBEC1. Interestingly, while Sdd6 showed nearly no off-target editing activity, it was still robust at on-target base editing when tested across six endogenous sites in rice protoplasts (Figure 4D and S4E). When we analyzed the on-target:off-target ratios of these ten deaminases, Sdd6 exhibited the highest on-target:off-target editing ratios, which was 37.6-fold that of rAPOBEC1 (Figure 4E). We further compared the on-target and off-target editing of Sdd6 to that of rAPOBEC1 and its two high-fidelity deaminase variants, YE1 and YEE, in HEK293T cells^43^. Importantly, we found that Sdd6 had the highest on-target:off-target editing ratios and was calculated to be 2.8-fold, 2.1-fold and 2.5-fold higher than that of rAPOBEC1, YE1 and YEE, respectively, and 10.4-fold higher than that of hA3A (Figure 4F and S4F). Notably, the on-target activity of Sdd6 was comparable to that of rAPOBEC1 and much higher than that of YE1 and YEE (Figure S4F). Thus, we identified that the SCP1.201 clade contains unique and more precise Sdd proteins to be used as high-fidelity base editors.

### Rational design of Sdd proteins assisted by AlphaFold2 structure prediction

Although viral delivery of CBEs has great potential for disease treatment, the large size of APOBEC/AID-like deaminases restricts their ability to be packaged into single AAV particles for *in vivo* editing applications^31^. Others have developed dual-AAV strategies delivery approaches by splitting CBEs into an amino-terminal and carboxy-terminal fragment and packaging them into separate AAV particles^31^. However, these delivery efforts would challenge large-scale manufacturing, require higher viral dosages, and pose potential safety challenges for human use^44^. Recently, a truncated sea lamprey cytidine deaminase-like 1 (PmCDA1)-based CBE was developed that could theoretically be packaged into a single-AAV, but the editing efficiency was extremely low when using the packaged AAVs during HEK293T cell transduction^45^. As SCP1.201 deaminases are canonically compact and conserved (Figure S5A), we thought that they might be the ideal protein for single-AAV CBEs.

We wondered whether we could use AI-assisted protein modeling to further engineer and shorten the size of the newly discovered Sdd proteins. We then generated multiple truncated variants of each of Sdd7, Sdd6, Sdd3, Sdd9, Sdd10, and Sdd4 and tested these variants for endogenous base editing in rice protoplasts across two sites each.

We identified mini-Sdd7, mini-Sdd6, mini-Sdd3, mini-Sdd9, mini-Sdd10, and mini-Sdd4 as newly minimized deaminases that are both small (∼130-160 aa) and have comparable or higher editing efficiencies compared to their full-length proteins both in rice protoplasts and human cells (Figure 5A, S5B and S5C). Strikingly, all six miniaturized deaminases would permit the construction of single-AAV-encapsulated SaCas9-based CBEs (< 4.7 kb between ITRs) (Figure 5B, S5D, S5E and S5F). We used mini-Sdd6 to construct a single-AAV SaCas9 vector and found that it had editing efficiencies of around 60% in mouse neuroblastoma N2a cells at two sites in the *HPD* gene (*Mus musculus* 4-hydroxyphenylpyruvate dioxygenase)^46^ by transient transfection (Figure 5C). These results highlight that the Sdd proteins offer great advantages over APOBEC/AID deaminases in terms of AAV-based CRISPR base editing delivery. The success in further shortening Sdd proteins for AAV packaging highlights the great potential of AI-assisted protein engineering.

**Figure 5.**
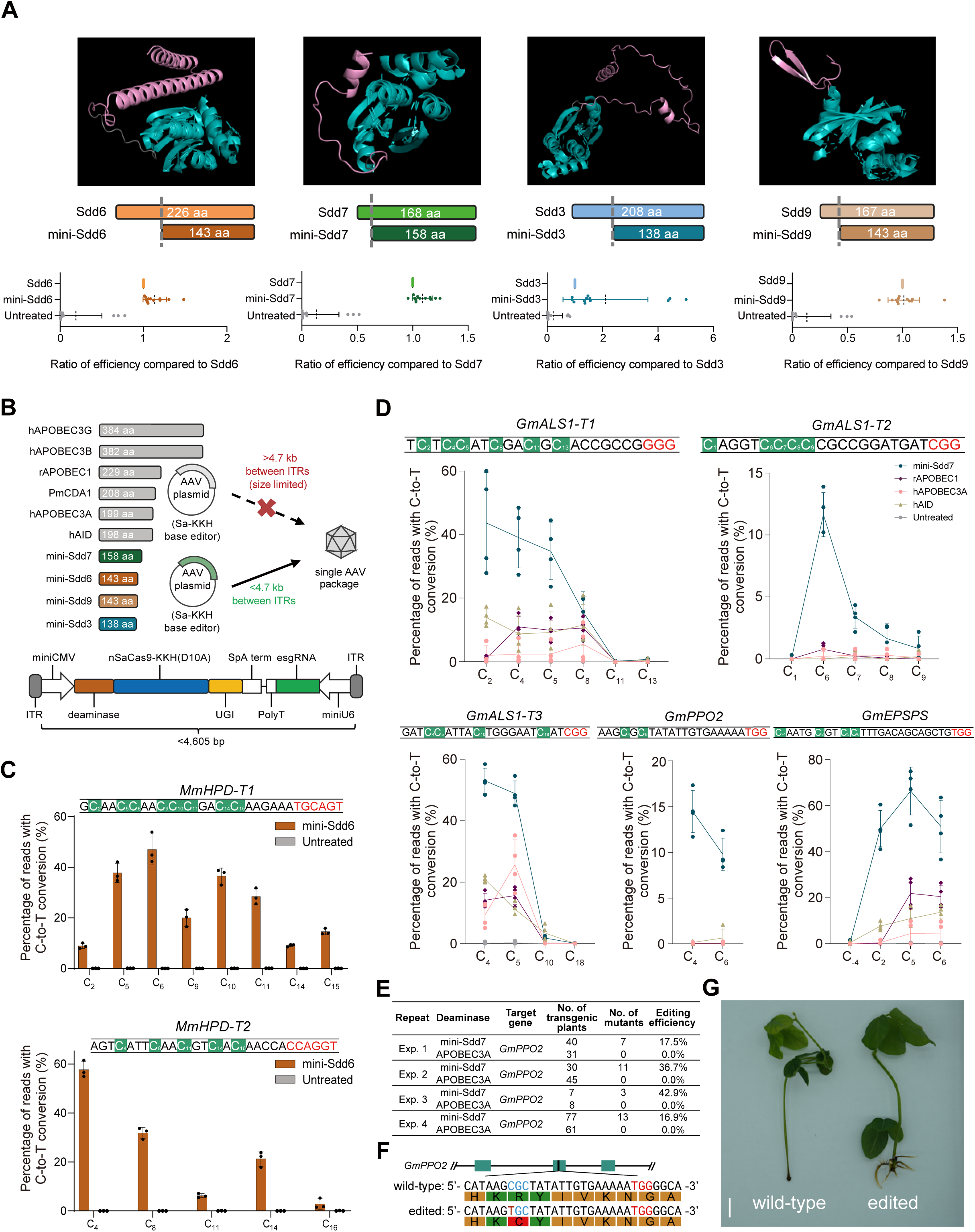
Engineering truncated Sdd proteins for use in animals and plants. (A) Engineering truncated Sdd proteins. Top panel, AlphaFold2-predicted structures of Sdd6, Sdd7, Sdd3, and Sdd9. Conserved regions are shown in cyan and truncated regions are shown in pink. Bottom panel, relative editing efficiencies of Sdds and their minimized version across two endogenous sites in rice protoplasts and two sites in HEK293T cells. (B) Theoretical packaging of a SaCas9-based CBE vector for packaging into a single AAV. Top panel, schematic diagram of APOBEC/AID-like deaminases, Sdds and their AAV vectors. Grayed deaminases are too large for single-AAV packaging. Bottom panel, schematic representation of Sdd-based AAV vectors. (C) Editing efficiency of mini-Sdd6 at two endogenous target sites in the *MmHP* gene in N2a cells. (D) Editing efficiencies of mini-Sdd7, rAPOBEC1, hA3A, and hAID base editors at five endogenous target sites in soybean hairy roots. (E) Frequencies of mutations induced by mini-Sdd7 and hA3A in T0 stable soybean plant editing in cotyledons by canonical *Agrobacterium tumefaciens*. The data were collected by four independent biological experiments. (F) The genotypes of base edited soybean plants. (G) Phenotypes of soybean plants treated with carfentrazone-ethyl for 10-days. Left panel, wild-type soybean plant (R98). Right panel, base-edited soybean plant (C98). Bar=1 cm. For (A), (C) and (D), Dots represent individual biological replicates, bars represent mean values, and error bars represent the s.d. of three or four independent biological replicates.

### Robust base editing with Sdd-based CBEs in rice and soybean

We next explored the use and application of newly engineered Sdd proteins for base editing in plants. We first evaluated the ability to use of mini-Sdd7 in *Agrobacterium*-mediated genome editing of rice plants and observed more mutants recovered and a greater proportion of edited plants, which reflects both a higher efficiency and lower toxicity compared to the most used hA3A-based CBE in agricultural application (Figure S5G).

Soybean is one of the most important staple crops grown around the world, serving as an essential source of vegetable oil and protein^47^. Although previously reported base editors have been widely used in many crops like rice, wheat, maize, potato and more, cytosine base editing remains challenging and poorly efficient across most sites tested in soybean crops^32, 48^. Since the first development of base editing, only one article has used *Agrobacterium tumefaciens* to obtain stable transformations and cytosine base-edited soybeans, but the efficiency was extremely low and resulted in chimeric plants rather than completely edited soybeans^32^.

We wondered whether our newly developed Sdd-based CBEs would result in superior cytosine base editing in soybeans. The transient base editing shown was evaluated using a soybean hairy root transformation mediated by *Agrobacterium rhizogenes*. This approach is often used in soybeans due to its quick nature (∼20 days) in allowing researchers to evaluate editing percentages in root cells. We constructed vectors with an AtU6 promoter driving sgRNA expression and a CaMV 2 × 35S promoter driving CBE expression and evaluated these using transgenic soybean hairy roots following *Agrobacterium rhizogenes*-mediated transformations (Figure S5H).

We found that the APOBEC/AID deaminases had low editing activities across all five sites evaluated as expected, including at the *GmALS1-T2* and *GmPPO2* sites which were particularly difficult to edit by other CBEs in soybean (Figure 5D). Remarkably, mini-Sdd7 displayed a 26.3-fold, 28.2-fold, and 10.8-fold increased cytosine base editing levels, respectively, compared to rAPOBEC1, hA3A and human activation-induced cytidine deaminase (hAID), respectively, across the five sites and reaching editing efficiencies up to 67.4% (Figure 5D). However, the cells from hairy root transformations are impossible to regenerate into soybean plants so the canonical *Agrobacterium tumefaciens* is used to perform stable soybean plant editing in cotyledons.

We next sought to use hA3A and mini-Sdd7 to base edit and obtain transgenic soybean plants following *Agrobacterium tumefaciens*-mediated transformation. We chose to edit the endogenous *GmPPO2* gene to create an R98C mutation, which would result in carfentrazone-ethyl resistant soybean plants^49^. Although the editing efficiencies from hairy root transformations are a great approach for evaluating relative editing efficiencies, it is not reflective of the percentage of edited plants following soybean plant regeneration. Even with the highly efficient hA3A-base editor in plants, we never successfully obtained cytosine base-edited plants (Figure 5E). Surprisingly, we obtained 34 base-edited heterozygotes from 154 transgenic soybean seedlings of Sdd7 transgenic plants from four independent biological experiments (Figure 5E). Therefore, Sdd7 now enables efficient cytosine base editing in soybean plants, which will greatly contribute to future agricultural breeding efforts (Figure 5E and 5F).

After treatment with carfentrazone-ethyl for ten days, we could obviously observe that while the wild-type plant was sensitive to wilting and could not generate roots, the mutated plant edited by Sdd7 grew well and normal (Figure 5G). The development of efficient cytosine base editors for use in soybean plants could enable diverse applications in the future.

## Discussion

Compared with the limited insights provided by 1D amino acid sequence alone, 3D structural information provides a more visually informative representation of potential protein functions. Structure-based protein mining promises to be a useful method for discovering and engineering new enzymes. Previously, research in functional genomics has been limited by either the cost of high-resolution analysis of protein structure or by the low-accuracy of traditional computational-driven folding simulations^50, 51^. AI-based high accuracy protein folding prediction models and the related databases have breathed new life into the life sciences.

Here we carried out a proof-of-concept exploration of protein classification and mining of novel protein functions based on structural predictions for the Cytidine Deaminase-like superfamily. We showed that AlphaFold2-predicted structures classified deaminases reliably into distinct clades with diverse protein folds and catalytic functions. We built on this by identifying deaminases with novel and different DNA substrates, which in turn permits the design of bespoke precision genome editing tools. In principle, this strategy could be applied to the high throughput classification and functional analysis of any protein dataset. We believe that future sequencing efforts in parallel with structural predictions will substantially advance the mining, tracking, classification, and design of functional proteins.

Currently only a few cytidine deaminases are in use as cytosine base editors. Canonical efforts based solely on protein engineering and directed evolution have helped diversify editing properties, however, these efforts are generally difficult to establish. Using our structure-based clustering methods, we discovered and profiled a suite of deaminases with distinct properties that can work both in plants and mammalian cells.

Among the newly AI rational discovered and designed deaminases, we identified compacted Sdd7 and Sdd6 to show great promise for both therapeutic and agricultural applications. Sdd7 was capable of robust base editing in all tested species and had much higher editing activity than the most commonly used APOBEC/AID-like deaminases. Surprisingly, we found that Sdd7 was capable of efficiently editing soybean plants, which was a major limitation for cytosine base editing previously. We speculated that Sdd7, derived from the bacterium *Actinosynnema mirum,* may possess high activity at temperatures suitable for soybean growth, in contrast to the mammalian APOBEC/AID deaminases. While profiling Sdd6, we found that this deaminase was smaller and by default more specific than the other deaminases while maintaining high on-target editing activity. We believe that these newer discovery and engineering efforts will contribute to the development of bespoke genome editing tools, which will be more precise and specific to each therapeutic or breeding application.

Advances in sequencing methods have propelled the discovery of new species and proteins. The advent of AI-assisted protein structure predictions in combination with growing numbers of sequencing efforts will further spark new enzyme discovery and enable even greater bioengineering efforts.

### Limitations of the study

Due to the length and time constraints of this paper, we cannot fully explore the properties of all proteins in the SCP1.201 family and other family proteins. However, we believe that in future studies, there will be many surprises for these large and unknown protein families.

## Acknowledgement

We thank Prof. Youwei Ai and Prof. Qingfeng Wu for kindly providing HEK293T and N2a cell lines, respectively. We thank Prof. Qi-jun Chen for kindly providing pBSE901. We thank Prof. Tianfu Han for kindly providing the seeds of Zhonghuang13 soybean.

This work was supported by the National Natural Science Foundation of China (32388201), the National Key Research and Development Program (2022YFF1002802), the Ministry of Agriculture and Rural Affairs of China, and the Strategic Priority Research Program of the Chinese Academy of Sciences (Precision Seed Design and Breeding, XDA24020102). Q.L is supported by Postdoctoral Innovative Talent Support Program of China (BX2021353) and China Postdoctoral Science Foundation (2022M720163). K.T.Z. was supported by the Schmidt Science Fellows.

## Author contributions

J.Huang, K.T.Z. and C.G. conceived the project and designed the experiments. J.Huang discovered the new deaminases. H.F. and Y.Li performed the structure-based protein classification analysis. J.Huang, Q.L., and Z.H. performed the protoplasts transformation and NGS data collection experiments. J.Huang, and Q.L. performed the mammalian cell transfection and NGS data collection experiments. J.Huang, H.F., and Z.H. collected the TRAP-seq data. J.Huang and Q.G. analyzed the Novaseq and Miseq data. G.L. and J.Hu prepared Miseq samples. H.F., and G.L. constructed the binary vectors for rice and soybean plant transformation. B.L. obtained regenerated rice plants. J.Huang, Z.H., and B.L. identified rice mutants. H.X., H.F., L.Z, Y.R., Z.H., and R.Z. performed soybean transformation, base-edited plants identification, and soybean resistance experiments. Y.Luo, K.Q., and P.H. generated the HEK293T cells with stable transfected TRAP-12K library. E.Z. provided AAV vector with guide RNA for mouse targets. Q.L. and Z.H. prepared the figures. C.G. and K.T.Z. and supervised the study. Q.L., H.F., Y.Li, K.T.Z. and C.G. wrote the manuscript with input from all authors. J.-L.Q. revised the manuscript.

## Declaration of interests

The authors have submitted two patent application based on the results reported in this paper. K.T.Z. is a founder and employee at Qi Biodesign.

## Inclusion and diversity

We support inclusive, diverse, and equitable conduct of research.

## STAR★Methods

### RESOURCE AVAILABILITY

#### Lead contact

Further information and requests for resources and reagents should be directed to and will be fulfilled by the Lead Contact: Caixia Gao.

#### Materials availability

All unique/stable reagents generated in this study are available from the Lead Contact with a completed Materials Transfer Agreement.

#### Data availability

The deep amplicon sequencing data were deposited in the PRJNA915939, PRJNA915940, PRJNA915941, and PRJNA915942. All other data are available in the main paper or supplement.

### EXPERIMENTAL MODEL AND SUBJECT DETAILS

#### *E.coli* transfection

FastT1 *E.coli* competent cells were used for amplifying plasmid DNA. Transfected *E.coli* cells were grown at 37°C in Lysogeny Broth (LB) medium supplemented with 100 mg/mL ampicillin or kanamycin overnight.

#### Rice protoplast transfection

For protoplasts transfection, we used the Japonica rice (*Oryza sativa*) variety Zhonghua11 to prepare protoplasts. Protoplast isolation and transformation were performed as described previously^52^. Plasmids (5 µg per construct) were introduced by PEG-mediated transfection. The transfected protoplasts were normally incubated at 26 ℃ for 72 hours for fluorescence cell observation or DNA extraction.

#### Mammalian Cell lines and culture conditions

Both human HEK293T cells (ATCC, CRL-3216) and mouse N2a cells (ATCC, CCL-131) were cultured in Dulbecco’s Modified Eagle’s medium (DMEM, Gibco) supplemented with 10% (vol/vol) fetal bovine serum (FBS, Gibco) and 1% (vol/vol) Penicillin-Streptomycin (Gibco) in a humidified incubator at 37 °C with 5% CO2.

## METHOD DETAILS

### Protein clustering and analyzing

Protein sequences were downloaded from InterPro database^53^ and NCBI’s BLAST^54^ (https://blast.ncbi.nlm.nih.gov/Blast.cgi) on the NR database. HMM was utilized to annotate deaminase domains to reduce the accumulation of unrelated information by HMMER^55^. We randomly chose 15 proteins from each family and clustered their domain sequences with a threshold of 90% sequence identity and 90% coverage using CD-HIT^56^. Representatives of each cluster were selected for further analysis. High confidence protein structures were predicted by Alphafold v2.2.0 and filtered with average per-residue confidence metric pLDDT ≥ 70.

Multiple sequence alignment was performed using Multiple Protein Sequence Alignment (MUSCLE)^57^. The phylogenetic tree was constructed using IQ-TREE 2 (http://www.iqtree.org) with 1500 ultrafast bootstraps^58^. A low perturbation strength (-pers 0.2) and large number of stop iterations (-nstop 500) were set because of the short length of the deaminase domains. Structure alignment was performed based on normalized TM-score^33^. The structural similarity matrix was further clustered by Unweighted Pair Group Method with Arithmetic mean (UPGMA) and visualized by Figtree (http://tree.bio.ed.ac.uk/software/figtree/). Protein structure diagrams were made in PyMOL^59^.

### Deaminase synthesis and removal of redundant sequence

We chose gene fragments encoding complete deaminase domains as well as extra N and C protein sequences for commercial synthesis (GenScript) (fig. S1). All of the candidate cytidine deaminases were codon optimized (rice and wheat or human and mouse). The toxin deaminase was split into two fragments and the split site was selected according to DddA by protein structure alignment. The conserved protein structure was obtained through multiple alignment of predicted structure in PyMOL^59^, which helps to conduct the removal of redundant sequence.

### Plasmid construction

For plant CBE vectors (maize ubiquitin-1 promoter-driven CBEs), synthesized deaminases were cloned into pnCas9-PBE vector (Addgene#98164), yielding vectors with Ubi-1::NLS-deaminase-linker-nCas9(D10A)-UGI-NLS::CaMV expression cassettes.

For CBE vectors for mammalian cells (CMV promoter-driven CBEs), synthesized deaminases-SpCas9-2UGI were cloned into p2T-CMV-ABEmax-BlastR vector (Addgene#152989), yielding vectors with CMV::NLS-deaminase-linker-nCas9(D10A)-2xUGI-NLS::bGH expression cassettes.

The DdCBE vectors including NLS, TALE array sequences, candidate cytidine deaminases, and UGI sequence were codon optimized for both human and mouse, synthesized commercially (Genscript), and cloned into pCMV_BE4max vector (Addgene#112093), yielding vectors with CMV::NLS-TALE-deaminase-UGI-NLS::bGH expression cassettes.

The plant sgRNA vectors (rice U3 promoter drives sgRNA) were constructed as reported previously using the pOsU3 backbone (Addgene#170132)^60^. To construct human and mouse sgRNA vectors (human U6 promoter drives sgRNA), the hU6 promoter was amplified and cloned into the pOsU3 backbone, followed by sgRNA target sequence cloning steps^52^.

Plant SaCas9 vectors for off-target testing were constructed as reported previously^42^.

To construct AAV vectors, the sequences between ITRs were synthesized (GenScript) and cloned into pX601 vector (Addgene#61591), followed by sgRNA target sequence cloning steps.

To construct binary vectors for rice plant transformation, the candidate cytidine deaminases were codon optimized, synthesized commercially (GenScript), and cloned into pH-nCas9-PBE vector (Addgene#98163), followed by sgRNA target sequence cloning steps^52^.

To construct binary vectors for soybean hairy root transformation, NLS, candidate cytidine deaminases, linker, nCas9(D10A), UGI, P2A, mScarlet sequences were codon optimized, synthesized commercially (GenScript), and cloned into pBSE901 (Addgene#91709) vector, followed by sgRNA target sequence cloning steps. To construct binary vectors for soybean transformation, the selection marker was replaced by the *EPSPS* sequence.

### Mammalian cell line transfection

All the cells were routinely tested for Mycoplasma contamination with a Mycoplasma Detection Kit (Transgen Biotech). The cells were seeded into 48-well Poly-D-Lysine-coated plates (Corning) in the absence of antibiotic. After 16-24 hours, cells were incubated with 1 µL Lipofectamine 2000 (ThermoFisher Scientific), 300 ng vector with deaminases, and 100 ng sgRNA expression vector. For DdCBEs transfection, cells were incubated with 1 µL Lipofectamine 2000, 300ng TALE-L and 300ng TALE-R. 72 hours later the cells were washed with PBS, followed by DNA extraction. For examining off-target effects by the R-loop assay, four vectors namely BE4max vector, SaCas9BE4max vector and the corresponding sgRNA vectors were co-transfected into cells^36^.

### TRAPseq library

We used the sgRNA 12K-TRAPseq library for evaluation of base editor properties. We seeded 2×10^6^ cells into 100 mm dish 20-hours before viral transduction. We transduced 500 µL of sgRNA lentivirus. For stably integrated cells, we used 1 µg/mL of puromycin (Gibco) to select. For each base editor, we seeded 2×10^6^ cells into 6-plates 24-hours before transfection. We transfected 15 µg of each CBE member plasmid DNA and 15 µg of Tol2 DNA with 60 µL of Lipofectamine 2000. Following 24 hours after transfection, we changed new culturing media to contain 10 µg/mL blasticidin (Gibco). After another 3 days, we washed the cells, suspended and reseeded all cells in 10 µg/mL blasticidin-containing media. After 6 days, we harvested all cells by washing with PBS then centrifuged and extracted DNA using Cell/Tissue DNA Isolation Mini Kit (Vazyme). For each member, we prepared sequencing reactions by applying 1.2 µg of DNA with a first set of primers following by barcoding and next-generating sequencing.

### DNA extraction

For HEK293T cells and N2a cells, genomic DNA was extracted with Lysis Buffer and Proteinase K with a Triumfi Mouse Tissue Direct PCR Kit (Beijing Genesand Biotech). For protoplasts, genomic DNA was extracted with a Plant Genomic DNA Kit (Tiangen Biotech) after 72 hours’ incubation. All DNA samples were quantified with a NanoDrop 2000 spectrophotometer (Thermo Scientific).

### Amplicon deep sequencing and data analysis

Triumfi Mouse Tissue Direct PCR Kit (Beijing Genesand Biotech) was used for amplification of target sequence in HEK293T cells and N2a cells. Phanta Max Master Mix (Vazyme) was used for amplification of target sequence in plants.

Nested PCR was used for amplification. In the first round PCR, the target region was amplified from genomic DNA with site-specific primers. In the second round, both forward and reverse barcodes were added to the ends of the PCR products for library construction. Equal amounts of PCR product were pooled and purified with a GeneJET Gel Extraction Kit (Thermo Scientific) and quantified with a NanoDrop 2000 spectrophotometer (Thermo Scientific). The purified products were sequenced commercially using the NovaSeq or Miseq platform, and the sequences around the target regions were examined for editing events^60^. Amplicon sequencing was repeated three times for each target site using genomic DNA extracted from three independent samples. Analysis of base editing behaviour by NovaSeq and Miseq was performed as described previously^60^.

For TRAP-seq analysis, we filtered NGS read depths of 12K TRAP below 50 and calculated the average editing efficiency at the corresponding surrogate target site inside the windows (from −10 to +27). In addition, we calculated the editing frequency for each NCN sequence motif and its proportions to evaluate context preferences.

### *Agrobacterium*-mediated transformation of rice calli

The Japonica rice (*Oryza sativa*) variety Zhonghua 11 was used for genetic transformation in this study. Binary vectors were introduced into *Agrobacterium tumefaciens* strain AGL1 by electroporation. *Agrobacterium*-mediated transformation of Zhonghua11 callus cells was conducted as reported^61^. Hygromycin (50 µg/ml) was used to select transgenic plants.

### Soybean hairy root transformation and plant transformation

The soybean (*Glycine max*) variety Williams 82 was used to generate hairy roots. Binary vectors were introduced into *Agrobacterium rhizogenes* strain K599 by electroporation. Explants were allowed to grow and develop roots for around 20 days in germination medium. Transgenic hairy roots were generated without selection in 10-12 days^62^. The soybean (*Glycine max*) variety Zhonghuang13 were used for generation of transgenic plants using *Agrobacterium tumefaciens*-mediated stable transformation. 10 mg/L glyphosate was used for selection during plant regeneration^63^. For phenotype identification of base-edited soybean, 0.3 mg/L carfentrazone-ethyl were added in rooting medium for selection.

### Plant mutant identification

Genomic DNA of transgenic plants was extracted with DNA Quick Plant System (Tiangen Biotech). Specific primers were used to amplify and sequence the target sites as described previously^60^ (Supplementary Table 1) (BGI). T0 transgenic rice and soybean plants were examined individually.

### Statistical analysis

All numerical values are presented as means ± s.d. Significant differences between controls and treatments were tested using the two-sided Mann-Whitney test, and *P* < 0.05 was considered statistically significant, *P* < 0.01 was considered statistically extremely significant.

## Supplemental information

### Primary Supplemental PDF

**Supplementary Table 1.** Primers used in this study.

