## Supplemental Figures for "Discovery of new deaminase functions by structure-based protein clustering"

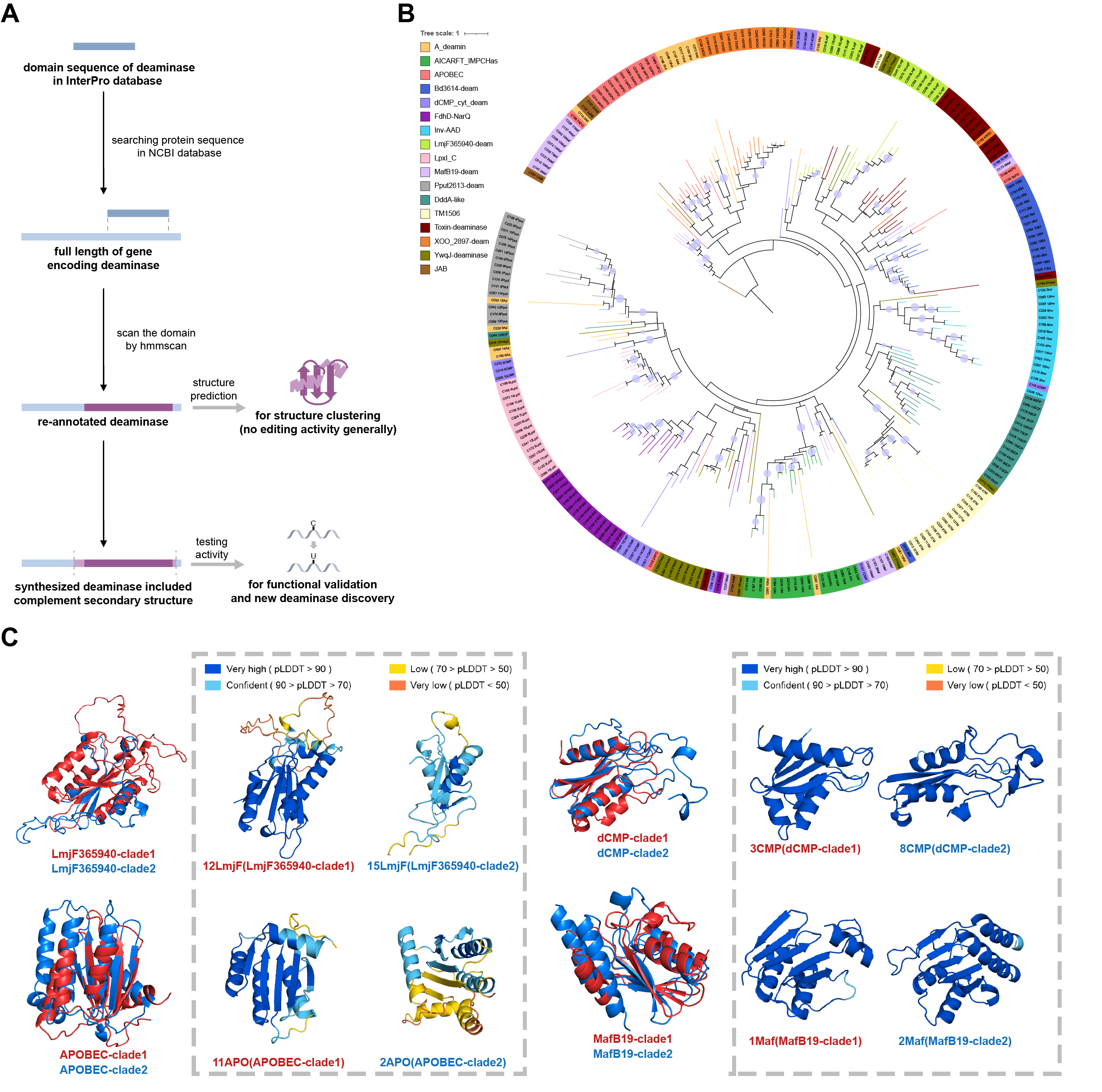

**Figure S1. Discovery of new cytidine deaminases via protein structures (related to Figure 1)**

(A) Workflow of re-annotation and synthesis of candidate deaminases. Since the amino acid sequence of the deaminase domains from InterPro may be incomplete, we used Protein BLAST from the NCBI database to obtain the full length of gene encoding deaminase, and then re-annotated the deaminase domain sequence with hmmscan (https://www.ebi.ac.uk/Tools/hmmer/search/hmmscan). The resulting domain sequences were then used for structure classification. Because the core deaminase domain used for clustering may not show editing activity, we synthesized some of the candidate deaminases with elongated N-terminal and C-terminal sequences from each corresponding gene. This extension will help to enhance protein stability and ensure the deaminase activity can be fully played, and then we evaluated their cytidine deaminase activity with the reporter system or at endogenous sites.

(B) Protein sequence-based inference of the phylogeny of the members of the CDA superfamily. Except for the JAB superfamily as outgroup, different cytidine deaminase families are shown by different color modes. Nodes with Bootstrap ≥ 90 are identified by circles.

(C) Alignment of representative structures of the separate LmjF365940, APOBEC, dCMP and MafB19 clades corresponding to Fig. 1b. The two represented structure of each clade was also showed with pLDDT. Although the pairs of clades from each of the four families had partially similar structures, their overall structures displayed relatively large differences, leading them to be classified as different.

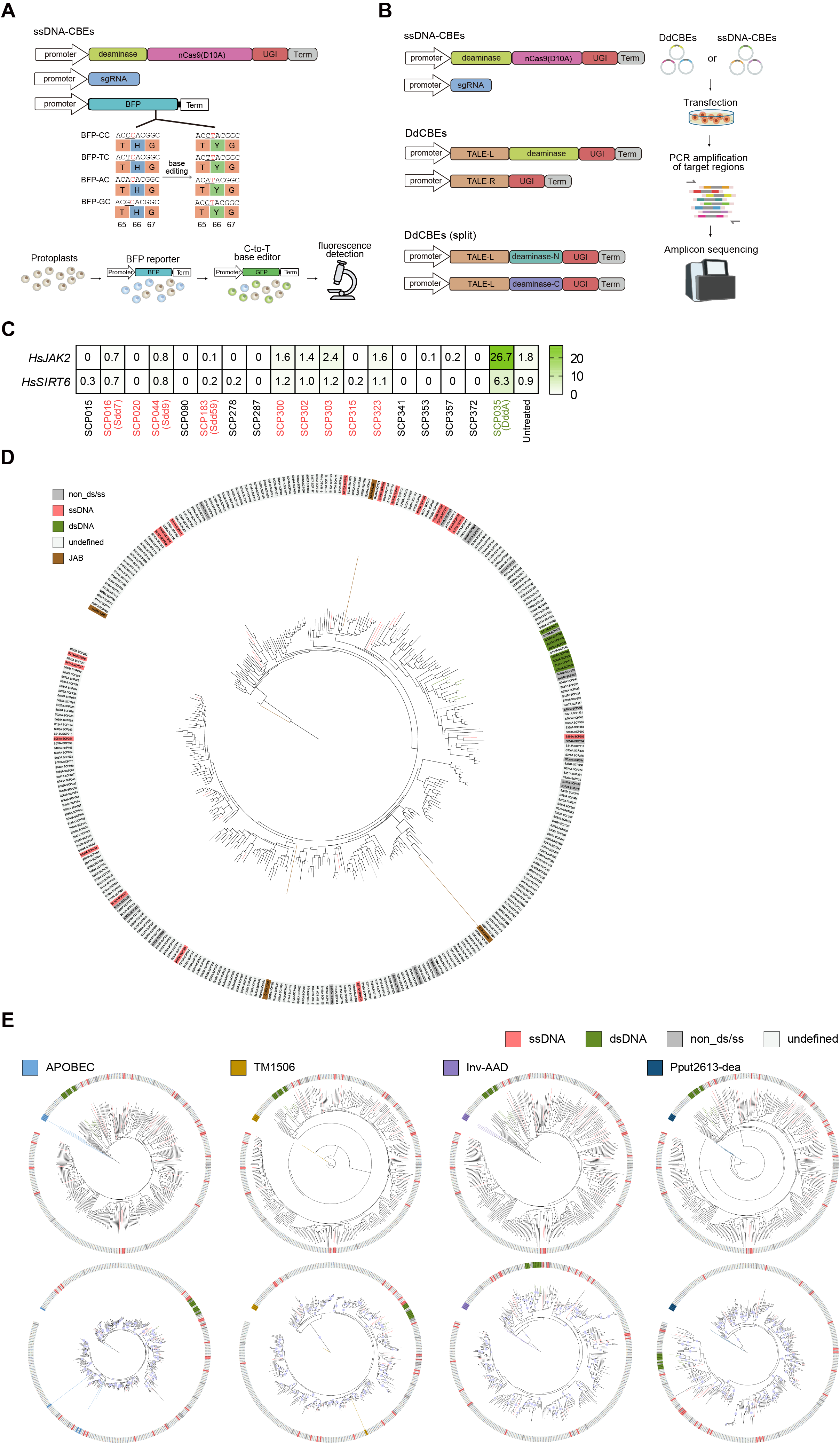

**Figure S2. Validation of the diverse functions of SCP1.201 deaminases (related to Figure 2)**

(A) Reporter system for ssDNA cytidine deamination activity identification. Top, schematic diagram of the ssDNA base editing vector for the BFP reporter system. Bottom, the procedure used to detect Sdd deaminases catalyzing C-to-T changes using the BFP reporter system in rice protoplasts.

(B) ssDNA and dsDNA cytidine deamination activity identification at endogenous sites. Left, schematic diagram of the ssDNA base editing vector for the endogenous site editing and the DdCBEs vector and its split form. Right. the procedure used to detect the activity of DdCBEs on dsDNA as well as ssDNA-CBEs on ssDNA in HEK293T cells, respectively, followed by high-throughput sequencing.

(C) Heatmap summarizing the editing efficiencies of dsDNA substrates of Sdd deaminases at *HsJAK2* and *HsSIRT6* sites. The gene name of active Sdd deaminase were colored as red. DddA with dsDNA deamination activity was colored as green. Data are representative of three independent experiments.

(D) Protein sequence-based inference of the phylogeny of the members of the SCP1.201 family. The JAB families were colored brown and regarded as an outgroup and the tested deaminases are shown in red, green and dark grey. Undefined deaminases in light grey await further functional analysis.

(E) Protein structure or sequence-based inference of the phylogeny of the members of the SCP1.201 family using four outgroups that more closely related to the SCP1.201 family. Top panel, based on protein structures; bottom panel, based on protein amino-acid sequences.

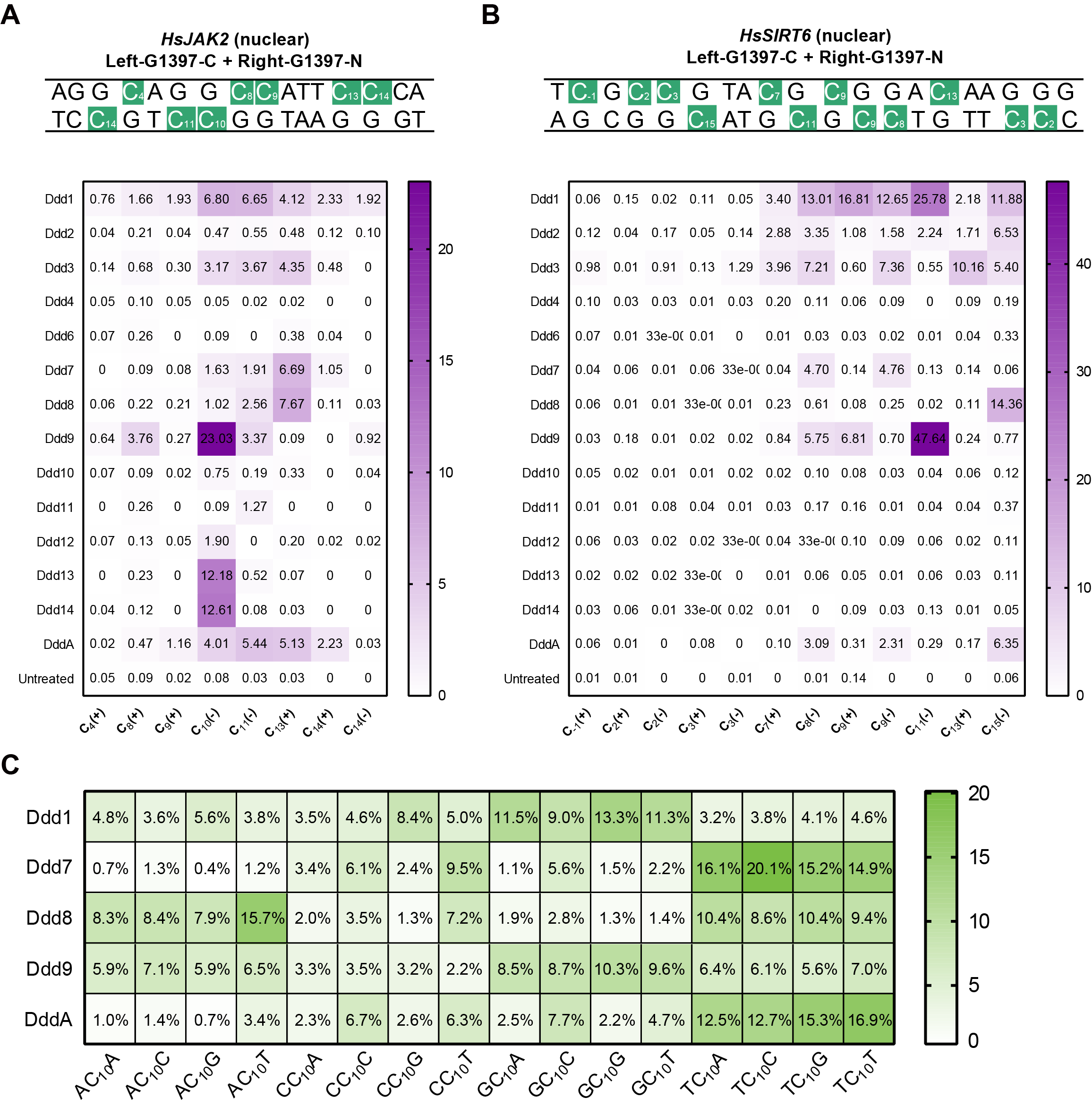

**Figure S3. Evaluating the activities and properties of newly discovered Ddd proteins for use as DdCBEs (related to Figure 3)**

(A) Heatmap of editing efficiencies and editing windows of SCP1.201 dsDNA deaminases at *HsJAK2* target sites in HEK293T cells.

(B) Heatmap of editing efficiencies and editing windows of SCP1.201 dsDNA deaminases at *HsSIRT6* target sites in HEK293T cells.

(C) The proportion of editing efficiencies of each context preferences among 16 plasmid libraries of different Ddd deaminases. Data are represented by the average of three independent experiments (*n* = 3).

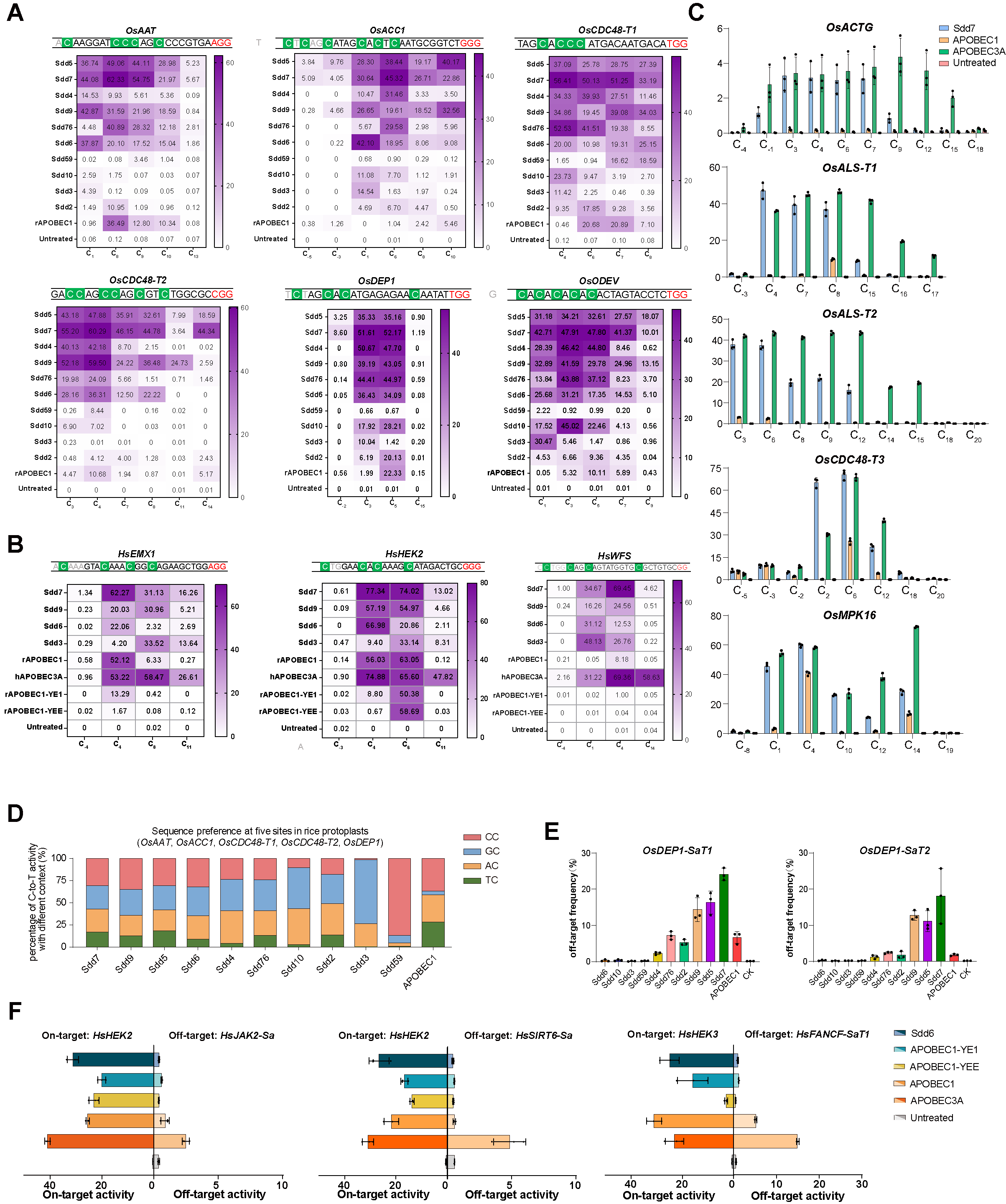

**Figure S4. Evaluating the activities and properties of newly discovered Sdd proteins for use as base editors (related to Figure 4)**

(A) Editing behavior of Sdd deaminases and APOBEC1 at six endogenous target sites in rice protoplasts. The heatmap shows the editing efficiencies and editing windows of ten Sdd deaminases and APOBEC1 at *OsAAT*, *OsACC1*, *OsCDC48-T1*, *OsCDC48-T2*, *OsDEP1* and *OsODEV* sites in rice protoplasts. The values given in the heatmap cells represents C-to-T editing efficiencies. Target sequences are listed above the heatmap, with green boxes marking the positions of C-to-T edits and PAMs in red font. Data are represented by the average of three independent experiments.

(B) Editing behavior of SCP1.201 ssDNA deaminases and APOBECs at three endogenous target sites in HEK293T cells. The heatmap gives the editing efficiencies and editing windows of four Sdd deaminases, rAPOBEC1, hA3A, rAPOBEC1-YE1, and rAPOBEC1-YEE at the *HsEMX1*, *HsHEK2*, *HsWFS1* sites in HEK293T cells. The values given in the heatmap cells represents C-to-T editing efficiencies. Target sequences are listed above the heatmap, with green boxes marking the positions of C-to-T edits and PAMs in red font. Data are represented by the average of three independent experiments.

(C) Comparison of the efficiencies of Sdd7, APOBEC1 and APOBEC3A at five sites in rice protoplasts. The efficiencies of Sdd7, APOBEC1 and APOBEC3A base editors compared across five endogenous targets, *OsACTG*, *OsALS-T1*, *OsALS-T2*, *OsCDC48-T3* and *OsMPK16*. Data are representative of three independent experiments.

(D) Sequence preference of Sdd deaminases and APOBEC1 at five endogenous target sites in rice protoplasts. The stacked graph shows the context preferences of ten Sdd deaminases and APOBEC1 at five endogenous target sites, *OsAAT*, *OsACC1*, *OsCDC48-T1*, *OsCDC48-T2* and *OsDEP1*. The green, yellow, blue and red bars represent the proportions of C-to-T activity for TC, AC, GC, and CC, respectively. Data are representative of three independent experiments.

(E) Frequencies of off-target events for Sdd deaminases and APOBEC1 at two endogenous target sites in rice protoplasts. Off-target events were evaluated using the orthogonal R-loop assay. Frequencies of off-target events for Sdd deaminases and APOBEC1 at the *OsDEP1-SaT1* and *OsDEP1-SaT2* sites in rice protoplasts. Data are representative of three independent experiments.

(F) On-target and off-target editing efficiency of Sdd6 and APOBEC base editors tested across two on-target and three off-target sites in HEK293T cells. Detailed display of on-target and off-target activity. On-target and off-target editing efficiencies of Sdd6, APOBEC1-YE1, APOBEC1-YEE, APOBEC1 and APOBEC3A across the *HsHEK2*, *HsHEK3* on-target site and the *HsJAK2-Sa*, *HsSIRT6-Sa* and *HsFANCF-SaT1* off-target sites. Data are representative of three independent experiments.

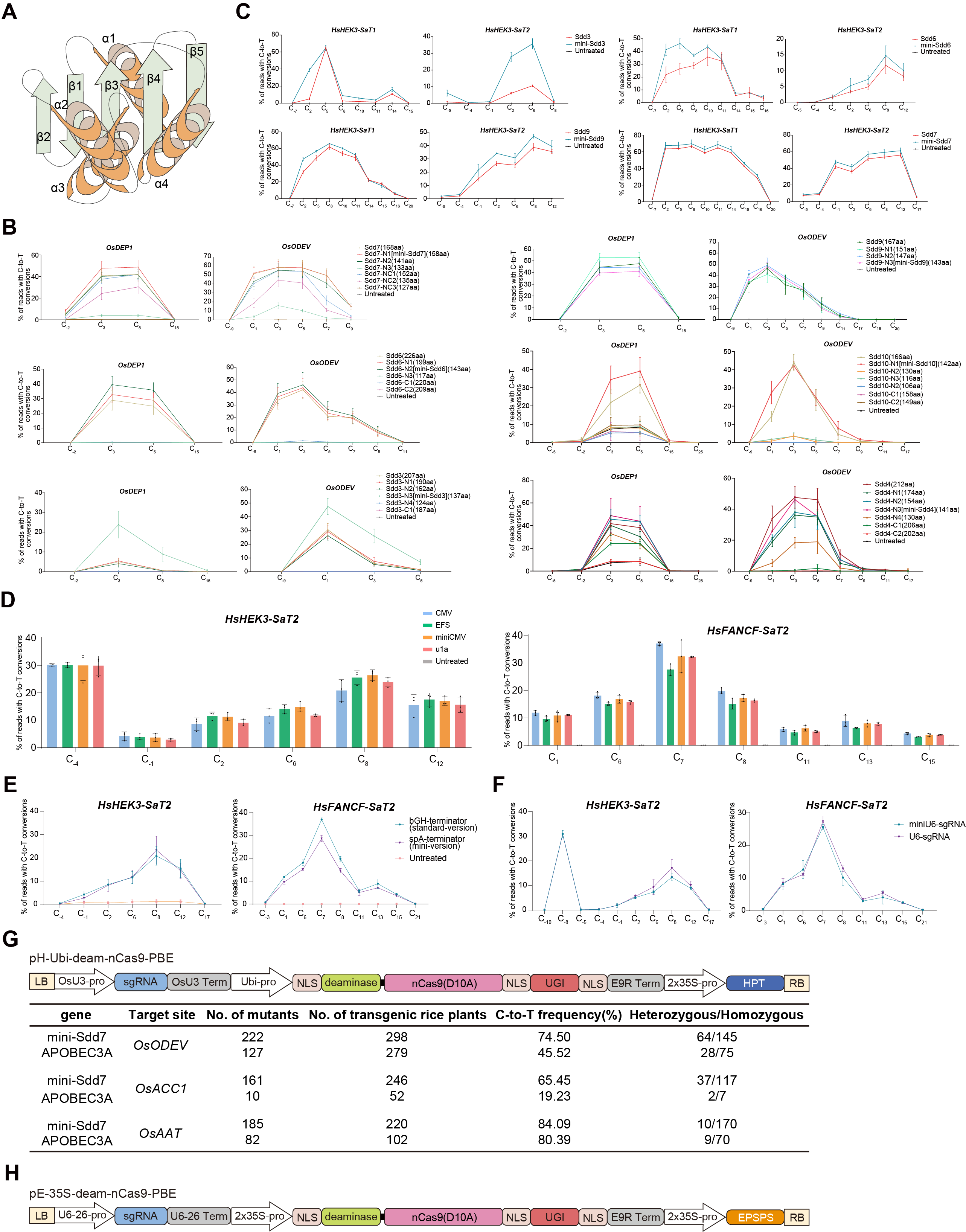

**Figure S5. Optimization and development of Sdd-CBEs for therapeutic and agricultural applications (related to Figure 5)**

(A) Conserved protein structure of Sdd deaminases with high activity predicted by AlphaFold2. The core structure of Sdd deaminases with high deamination activity is shown. For some deaminases, α4 is not essential.

(B) Testing of the efficiencies of different truncated versions of synthesized deaminase genes at two endogenous target sites in rice protoplasts. Removal of redundant sequence of synthesized deaminase genes assisted by AlphaFold2. Editing efficiencies of multiple redundant sequence removal versions of Sdd7, Sdd6, Sdd3, Sdd9, Sdd10 and Sdd4 deaminases at the *OsDEP1* and *OsODEV* sites in rice protoplasts are showed. Truncations included various forms of C-terminal and N-terminal deletions. Data are representative of three independent experiments.

(C) Testing of the efficiencies of different truncated versions of synthesized deaminase genes at two endogenous target sites in human cells. Removal of redundant sequence of synthesized deaminase genes assisted by AlphaFold2. Comparison of the Sdd3, Sdd9, Sdd6, Sdd7 deaminases and their redundant sequence removal versions at the *HsHEK3-SaT1* and *HsHEK3-SaT2* sites in HEK293T cells. Data are representative of three independent experiments.

(D) Comparison the editing efficiencies of mini-Sdd6 with four promoters, CMV, EFS, mini-CMV, and u1a at the *HsHEK3-SaT2* and *HsFANCF-SaT2* sites in HEK293T cells.

(E) Comparison of mini-Sdd6 with two terminators, bGH and SpA, at the *HsHEK3-SaT2* and *HsFANCF-SaT2* sites in HEK293T cells, to observe the effects of terminators on AAV vectors.

(F) Comparison of minU6 and U6 promoters at the *HsHEK3-SaT2* and *HsFANCF-SaT2* sites in HEK293T cells. Data are representative of three independent experiments.

(G) Frequencies of base edited re-generated rice plants. Top, schematic diagram of the base editing binary vector for *Agrobacterium*-mediated transformation in rice. Bottom, frequencies of mutations induced by mini-Sdd7 and APOBEC3A base editors in T_0_ rice plants.

(H) Schematic diagram of the base editing binary vector for *Agrobacterium*-mediated transformation in soybean.
